# Influence of non-content instructor talk on students’ motivation-related outcomes in laboratory courses

**DOI:** 10.64898/2026.05.13.724928

**Authors:** Christopher James Zajic, Erin L. Dolan

## Abstract

Course-based undergraduate research experiences (CUREs) can expand undergraduates’ access to research and motivate students to stay in science. Yet, little research has examined how CURE instruction shapes student motivation. We leveraged a motivation-related characterization of non-content talk of 48 CURE and non-CURE instructors to predict the motivation-related outcomes of 462 students. We fit a series of multi-level models (MLM) in which we regressed students’ post-course scientific self-efficacy, task values, scientific identity, and science-related intentions onto instructors’ self-efficacy and task values-related talk, controlling for students’ pre-course levels. We also fit an MLM to explore whether instructors’ relationship-building talk (immediacy talk) was associated with students’ rapport with their instructor. Instructors’ self-efficacy talk did not affect students’ self-efficacy, and instructors’ immediacy talk had a marginally positive but non-significant association with students’ rapport ratings. Instructors’ task values talk positively influenced students’ scientific identity and some but not all of their task values. Instructors’ task values talk also positively influenced students’ intentions to pursue a science career, but not graduate education or research careers. Collectively, these results suggest that instructors’ task values talk may underpin some of the motivational effects of CURE instruction, but that task values talk need not be limited to CUREs.

**HIGHLIGHT:** We examine whether instructor talk predicts students’ motivational outcomes in CURE and non-CURE lab courses. Self-efficacy talk had no effect on student self-efficacy. Task values talk positively affected students’ science identity and career intentions, and some value beliefs. Immediacy talk was marginally related to student-instructor rapport.

## INTRODUCTION

Course-based undergraduate research experiences (CUREs) have been championed for their potential to engage many more life science students in undergraduate research than is possible through apprenticeship-style internships (Auchincloss et al., 2014). Advocacy for CUREs has led to their widespread development and implementation within and beyond the life sciences (Buchanan & Fisher, 2022; Watts & Rodriguez, 2023). Several studies have indicated that completing a CURE in place of a non-research laboratory course can influence students’ persistence in STEM majors and completion of college degrees (Hanauer et al., 2017; Jordan et al., 2014; Rodenbusch et al., 2016). These studies have examined only a handful of CUREs that span multiple semesters and have not offered insights into the aspects of CURE instruction that lead to these outcomes.

A decade ago, two papers presented theories about the elements of CUREs that, when combined, would make them distinctive and influential for students when compared with traditional, non-research lab courses (Auchincloss et al., 2014; Brownell & Kloser, 2015). Specifically, CURE students are thought to engage in multiple scientific practices; have opportunities to make broadly relevant discoveries; engage in iterative work such as troubleshooting, problem-solving, and repeating or replicating experiments; and collaborate in ways that are interdependent (i.e., relying on one another, building on one another’s work). Some research has supported these hypotheses, linking one or more of these design elements to indicators of students’ intentions to continue in science (Cooper et al., 2019; Corwin et al., 2015, 2018). Another line of inquiry has explored project ownership as a design feature that makes CUREs influential for students (Hanauer et al., 2012, 2017; Hanauer & Dolan, 2014). However, research connecting CURE design and implementation to student outcomes indicates that opportunities to make relevant discoveries, engage in iterative work, and develop a sense of ownership of coursework explains only a modest portion of the variation in students’ intentions to continue in science from pre to post CURE (Corwin et al., 2018). Collectively, this research indicates there are other aspects of CURE curriculum and instruction that are likely to be influential for students and as yet unidentified.

## THEORETICAL FRAMEWORK

Theories of motivation may be useful for gaining novel, mechanistic insight into *how* CURE instruction leads to students’ continuation in college and in science. Specifically, situated expectancy value theory (SEVT) and its predecessor, expectancy value theory (EVT), have been leveraged to explain students’ motivated behavior in academic and career decision-making settings (Eccles & Wigfield, 2002, 2020). Core tenets of this theory are that an individual’s expectations of success on a task and their beliefs about the value of the task shape their decisions to engage in the task. When applied to undergraduate science majors, this theory predicts that students’ beliefs about their ability to be successful in science (i.e., their scientific self-efficacy) and their beliefs about the value of doing science (i.e., their subjective task values) shape whether they decide to continue in science (i.e., motivated behavior). Some research has shown that CURE instruction can foster growth in students’ scientific self-efficacy (see Dolan, 2016 for review; Liu, 2022; Martin et al., 2021; Mendez et al., 2025) and influence students’ science task values (Ceyhan & Tillotson, 2020). Furthermore, an individual’s perceptions of the beliefs and behaviors of “socializers” in their environment, such as the instructors who teach their courses, influence their competency and value beliefs (Eccles & Wigfield, 2020). However, little research has aimed to causally link aspects of CURE design and instruction to changes in students’ expectancies and value beliefs. This kind of research is necessary to inform efforts to design and teach CUREs and other forms of lab instruction in a way that aligns with goals for students.

Research from Zajic and colleagues (2026) used EVT as a frame to characterize and compare non-content instructor talk, meaning the things instructors say beyond the content of the course (Seidel et al., 2015), in CUREs and non-CUREs. This work described three main forms of non-content instructor talk that students might perceive as motivating (Zajic et al., 2026). First, instructors used “self-efficacy talk,” meaning things instructors said that could influence students’ beliefs that they could be successful in science. One example of self-efficacy talk was normalizing the struggles of doing science, which could assuage students’ concerns that scientific failures are their fault. Another example of self-efficacy talk was providing encouragement, including complimenting students’ work, which could bolster students’ beliefs that they are capable of being successful in science. Second, instructors used “task values talk,” meaning things instructors said that could influence students’ beliefs that doing science offers sufficient benefits considering potential costs. One example was fostering wonder, initially described by Harrison and colleagues (Harrison et al., 2019), which is when an instructor expresses interest in or enthusiasm about students’ scientific work, which could in turn spark students’ interest or enthusiasm. Another example is articulating broad relevance, which is when an instructor comments on the larger purpose of students’ scientific work, possibly enhancing students’ beliefs that their work is valuable and recognizing their role in the scientific community (i.e., science identity; (Carlone & Johnson, 2007; Chemers et al., 2011; Estrada et al., 2011; Pfeifer et al., 2024)). Finally, instructors used “immediacy talk,” meaning things instructors said that could promote students’ sense of closeness with and positive feelings about the instructor (Mehrabian, 1966; Wiener & Mehrabian, 1968), in turn motivating student engagement in coursework and ultimately their learning and performance (Christophel, 1990; Christophel & Gorham, 1995; Gorham, 1988). Notably, this characterization of instructor talk from a motivational perspective revealed some differences in the types of talk used by CURE and non-CURE instructors, but also substantial overlap, primarily because of wide variation in the talk used by CURE instructors.

Results from Zajic and colleagues (2026) complement findings from other studies of CURE instruction, all of which show that the design and implementation of these courses vary widely and overlap in meaningful ways with other forms of lab instruction (Beck et al., 2023; Watts & Rodriguez, 2023; Zajic et al., 2026). Collectively, this research points to the importance of moving away from studying CUREs as a singular experience and toward identifying the instructional variables at play in CUREs or other forms of laboratory education. Furthermore, this prior work compared talk between course types but did not test whether particular types of talk were predictive of students’ motivation-related outcomes (Zajic et al., 2026). A critical next step is to test the relationships between types of instructor talk (i.e., self-efficacy, task values, and immediacy talk) and the student outcomes they are predicted to influence.

### CURRENT STUDY

The current study aims to determine whether instructor talk type predicts students’ motivation-related outcomes. Specifically, we surveyed students in 24 CURE and 24 non-CURE courses at the start and end of their course about their scientific self-efficacy, beliefs about the benefits and costs of doing research (i.e., task values), scientific identity, and intentions to continue in science-related education and career paths. We also surveyed students at the end of their course about their perceived rapport with their instructor. Then we leveraged an EVT-driven characterization of non-content instructor talk in these courses (Zajic et al., 2026) to fit a series of multilevel regression models to address the following research questions:

1. To what extent does instructor self-efficacy talk influence students’ post-course scientific self-efficacy beliefs, controlling for pre-course levels?
2. To what extent does instructor task values talk influence students’ post-course task value beliefs and scientific identity, controlling for pre-course levels?
3. To what extent is instructor immediacy talk associated with student ratings of rapport with their instructor?
4. To what extent do self-efficacy, task values, and immediacy talk influence students’ intentions to continue in science?

## METHODS

To address our research questions, we fit multi-level linear models (MLMs) to prior qualitative content analysis of non-content instructor talk (Zajic et al., 2026) to predict post-course student outcomes, controlling for pre-course values and accounting for nesting of students within courses. The results reported here are part of a larger study that was reviewed and determined to be exempt by the University of Georgia Institutional Review Board (PROJECT00003103).

### Recruitment, Participants, and Data Collection

Our recruitment methods and study sample are detailed in Zajic et al. (2026) and described briefly here for context. We recruited instructors of introductory life science laboratory courses (i.e., practical courses that involved hands-on bench, field, or computational work), from colleges and universities across the U.S. and Canada via direct emails and listerv announcements. Ultimately, 48 instructors, 24 who taught CUREs and 24 who taught non-CUREs, submitted audio recordings of a subset of their lab course sessions, and thus were included in our final analytic sample (Table 1). We asked participating instructors to share a study announcement with their students, which included a link to the consent information and the pre-course survey. We emailed students who completed the pre-course survey again at the end of their course to ask them to complete the post-course survey. In total, 882 students completed the pre-survey and 609 responded to the post-survey. We removed 147 students due to incomplete responses, resulting in a final analytic sample of 462 students (Table 2).

**Table 1.**
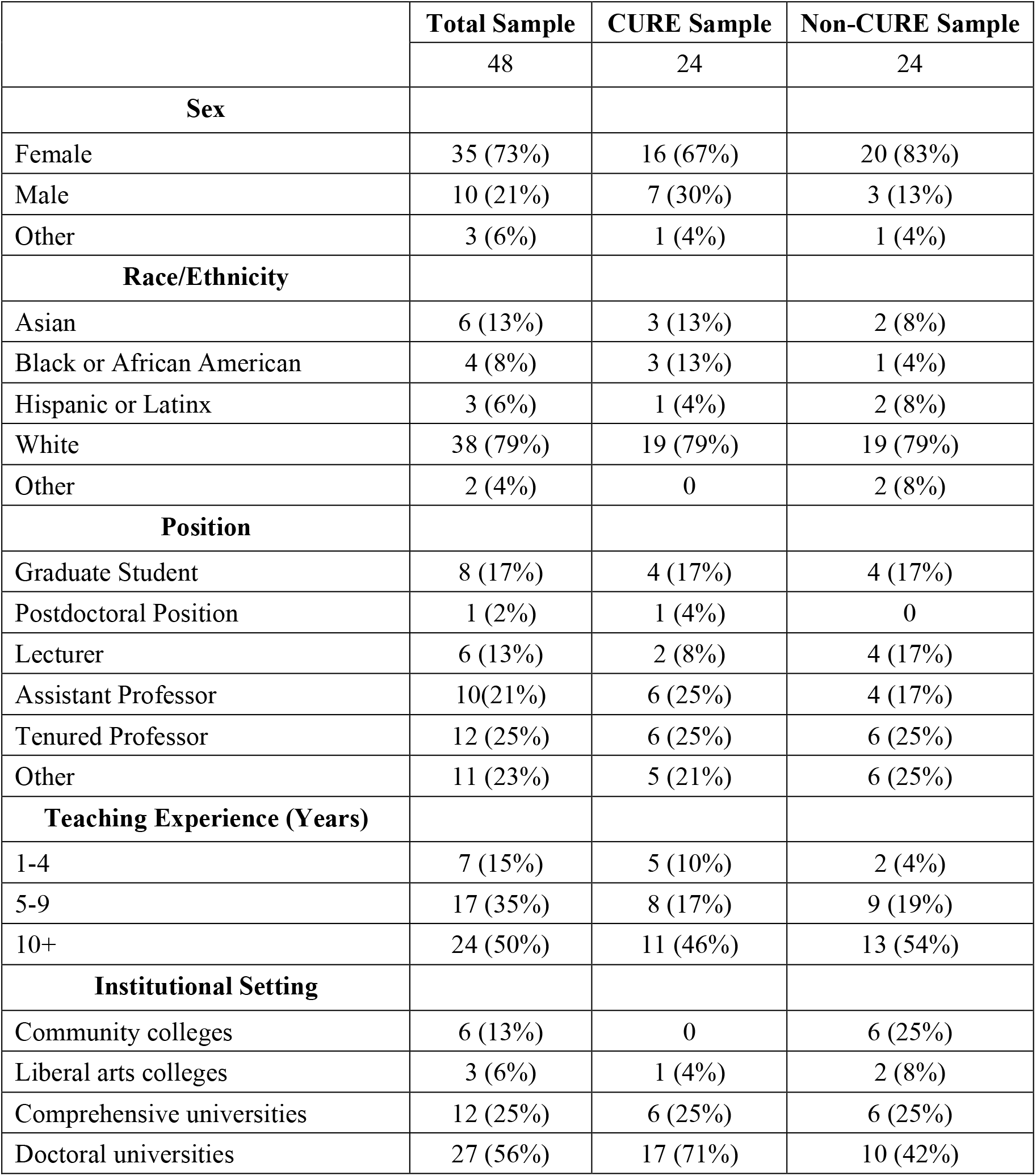
Instructor demographics. Percentages reflect proportion within the total sample or each sub-sample (i.e., within CUREs or non-CUREs). Originally published in Zajic et al. (2026) and included here for context.

**Table 2.** Student demographics. Percentages reflect proportion within the total sample or each sub-sample (i.e., within CUREs or non-CUREs). Students had the option to select multiple races/ethnicities. Thus, percentages within these demographics may total to >100%. Originally published in Zajic et al. (2026) and included here for context.

|  | **Total Sample** | **CURE Sample** | **Non-CURE Sample** |
| --- | --- | --- | --- |
|  | 462 | 261 | 201 |
| **Sex** |  |  |  |
| Female | 349 (76%) | 201 (77%) | 148 (74%) |
| Male | 91 (20%) | 49 (19%) | 42 (21%) |
| Other | 10 (2%) | 6 (2%) | 4 (2%) |
| Prefer not to respond | 12 (3%) | 5 (2%) | 7 (3%) |
| **Race/Ethnicity** |  |  |  |
| American Indian or Alaskan Native | 4 (<1%) | 2 (<1%) | 2 (<1%) |
| Asian | 130 (28%) | 80 (31%) | 50 (25%) |
| Black or African American | 49 (11%) | 31 (12%) | 18 (9%) |
| Hispanic or Latinx | 76 (16%) | 52 (20%) | 24 (12%) |
| Native Hawaiian or Other Pacific Islander | 2 (<1%) | 2 (<1%) | 0 |
| White | 235 (51%) | 121 (46%) | 114 (57%) |
| Other | 11 (2%) | 7 (3%) | 4 (2%) |
| Prefer not to respond | 8 (2%) | 3 (1%) | 5 (2%) |
| **Year in College** |  |  |  |
| First Year | 262 (57%) | 177 (68%) | 85 (42%) |
| Second Year | 115 (25%) | 49 (19%) | 66 (33%) |
| Third Year | 46 (10%) | 18 (7%) | 28 (14%) |
| Fourth Year | 28 (6%) | 17 (7%) | 11 (5%) |
| Other | 8 (2%) | 0 | 8 (4%) |
| Prefer not to respond | 3 (<1%) | 0 | 3 (1%) |
| **Prior Research** |  |  |  |
| None | 173 (37%) | 107 (41%) | 66 (33%) |
| 1 Term | 146 (32%) | 85 (33%) | 61 (30%) |
| 2 Terms | 76 (16%) | 44 (17%) | 32 (16%) |
| 3 Terms | 26 (6%) | 9 (3%) | 17 (8%) |
| >3 Terms | 32 (7%) | 13 (5%) | 19 (10%) |
| Prefer not to respond | 9 (2%) | 3 (1%) | 6 (3%) |

### Data Sources and Measures

We used two main data sources for this study: qualitative characterization of instructor talk and student survey responses. The methods used to characterize instructor talk are detailed in Zajic et al. (2026) and described briefly here for context. We used qualitative thematic analysis to identify instances of non-content instructor talk using an expectancy value theory frame (Zajic et al., 2026). In this prior work, we identified 716 instances of immediacy talk (35% of the corpus), 683 instances of self-efficacy talk (34% of the corpus), and 627 instances of task values talk (31% of the corpus). We summed all instances of immediacy talk, self-efficacy talk, and task values talk for each instructor, divided this value by the total minutes of recording for that instructor, and multiplied by 60 to generate three values for each instructor: immediacy talk per hour, self-efficacy talk per hour, and task values talk per hour. We used these values as the instructor talk data in the current study.

To examine the effects of instructor talk on students, we surveyed students at the start and end of their lab courses using the secure survey platform, Qualtrics™. We used the following established measures of our outcomes of interest:

- *Scientific Self-Efficacy*. We used a 9-item scale (Chemers et al., 2011; Estrada et al., 2011) to measure students’ confidence in their ability to conduct various science research practices (i.e., their scientific self-efficacy). A sample item is, “Please indicate how confident you are in your ability to trouble-shoot an investigation or experiment.” Response options ranged from 1 (not confident) to 6 (very confident).
  ∘ *Task Values* (Gaspard et al., 2015): Response options for all ranged from 1 (strongly disagree) to 6 (strongly agree).
  ∘ *Intrinsic Value*. We used a 3-item subscale to measure students’ personal interest and enjoyment (i.e., intrinsic value) associated with doing research. A sample item is, “Research is fun to me.”
  ∘ *Attainment Value*. We used a 3-item subscale to measure students’ beliefs about the importance of doing well (i.e., attainment value) in research. A sample item is, “Performing well in research is important to me.”
  ∘ *Social Utility*. We adapted a 3-item subscale to measure students’ beliefs about the social or communal value doing research. A sample item is, “I can do good in the world based on my knowledge of research.”
  ∘ *Job Utility*. We used a 3-item subscale to measure students’ beliefs about the utility of doing research for their future careers. A sample item is, “Learning how to conduct research is worthwhile because it improves my career prospects.”
  ∘ *Life Utility*. We used a 3-item subscale to measure students’ beliefs about the utility of doing research in their daily lives. A sample item is, “I will often need research in my life.”
  ∘ *Opportunity Costs*. We used a 3-item scale to measure students’ beliefs about the opportunity costs of doing research. A sample item is, “I’d have to sacrifice a lot of free time to be successful at research.”
- *Science Identity*. We used a 7-item scale (Chemers et al., 2011; Estrada et al., 2011) to measure the extent to which students identified as scientists. A sample item is, “I think of myself as a scientist.” The response options ranged from 1 (strongly disagree) to 6 (strongly agree).
- *Intentions*. We adapted 3 individual items (Estrada et al., 2011) to measure students’ intentions to pursue graduate school, a career in science, and a career in science research. Response options ranged from 1 (I definitely will not pursue a…) to 5 (I definitely will pursue a…).

We also included an 11-item scale (Frisby & Martin, 2010) on the post-survey to measure students’ perceptions of their *rapport* with their instructor. We opted not to measure rapport at the beginning of the course because we assumed students were unlikely to know their instructor and would not have a basis for judging their rapport. A sample item is, “I enjoy interacting with my instructor.” The response options ranged from 1 (strongly disagree) to 7 (strongly agree).

To assess the internal structure of each measure, we conducted confirmatory factor analyses using the “lavaan” package (Rosseel, 2012) in R statistical software (R Core Team, 2022). See Supplemental Materials for full descriptions of each scale and measurement model fit.

## Data Analysis

We calculated descriptive statistics for all student constructs of interest (e.g., means, standard deviations; see Supplemental Materials for correlations). We also conducted paired t-tests to identify pre- to post-course changes for all student variables except ratings of instructor rapport, which was only measured post-course. We employed multi-level linear models (MLMs) to determine the extent to which instructor talk predicted student outcomes, accounting for nesting of students within courses. For all MLMs, we utilized the “lme4” package (Bates et al., 2015) in R statistical software (R Core Team, 2022). We calculated the within and between-group variances (i.e., the intraclass correlations) for each student outcome variable (see Supplemental Materials). We fit a total of 18 MLMs to test our theorized relationships between instructor talk and student outcomes. We group-mean centered our level 1 predictors (i.e., students’ pre-course scores) and grand-mean centered our level 2 predictors (i.e., instructor talk), as has been previously recommended (Enders & Tofighi, 2007). The general structure of our models took on the following form:

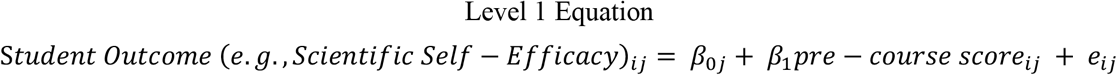

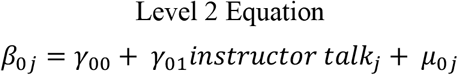

In the above equations, *β*_0j_ represents the mean pre-course score, or the intercept, for students within a given cluster, or instructor, *j. β*_1_ is the regression coefficient, or the fixed effect, of students’ pre-course scores on their post-course scores. *e*_ij_ represents the residual error for student *i* within instructor *j*’s class. *γ*_00_ represents the mean intercept for all instructors’ use of instructor talk. *γ*_01_ is the regression coefficient, or the fixed effect, associated with instructors’ use of instructor talk. *μ*_0j_ is the residual error at the instructor level.

## RESULTS

### Self-efficacy talk does not influence students’ scientific self-efficacy

We hypothesized that instructors’ self-efficacy talk would boost their students’ scientific self-efficacy from pre- to post-course. To test this hypothesis and address our first research question (RQ1: *To what extent does instructor self-efficacy talk influence students’ post-course scientific self-efficacy beliefs, controlling for pre-course levels?*), we conducted a paired t-test to compare students’ pre- and post-course self-efficacy. Consistent with prior research on CUREs (e.g., Liu, 2022; Martin et al., 2021; Mendez et al., 2025), students reported significant increases in their scientific self-efficacy from pre- to post-course (Table 3). We then regressed students’ post-course scientific self-efficacy onto the frequency of their instructor’s self-efficacy talk. We found that instructors’ self-efficacy talk had no effect on students’ self-efficacy (Table 4). This result suggests that the things instructors say that have the potential to build students’ confidence, such as noting that experimental failures are not students’ fault or complimenting students’ work (Zajic et al., 2026), are not having a detectable effect on students’ confidence in their ability to do science. Furthermore, the absence of a detectable effect of instructors’ self-efficacy talk is not due to a ceiling effect, given the observed increase in students’ self-efficacy.

**Table 3.** Descriptive statistics and pre- to post-course comparisons of student outcomes. This table presents means and standard deviations for all student variables at the start (pre) and end (post) of their course. The results of paired t-tests are also presented; **\* bolded font** indicates significant differences at *p* < 0.001. [**Note:** Rapport is on a 7-point scale, intentions are on a 5-point scale, and all other variables are on a 6-point scale.]

| Variable |  | Pre |  | Post |  | Mean Diff<br>[95% CI] | df | t | p |
| --- | --- | --- | --- | --- | --- | --- | --- | --- | --- |
|  |  | Mean | SD | Mean | SD |  |  |  |  |
| **Scientific self-efficacy \*** |  | **3.58** | **0.81** | **4.19** | **0.94** | **0.61**<br>**[0.53, 0.70]** | **460** | **14.1** | **<0.001** |
| Task Values | Intrinsic Value | 4.33 | 1.23 | 4.40 | 1.36 | 0.07<br>[-0.03, 0.17] | 460 | 1.38 | 0.17 |
|  | Attainment Value | 4.78 | 1.12 | 4.70 | 1.28 | -0.08<br>[-0.18, 0.02] | 460 | -1.58 | 0.12 |
|  | Social Utility | 4.81 | 1.00 | 4.74 | 1.07 | -0.08<br>[-0.17, 0.01] | 459 | -1.69 | 0.09 |
|  | **Job Utility \*** | **5.18** | **0.91** | **5.02** | **1.02** | **-0.17**<br>**[-0.25, -0.08]** | **460** | **-3.78** | **<0.001** |
|  | Life Utility | 4.55 | 1.03 | 4.51 | 1.16 | -0.04<br>[-0.14, 0.05] | 459 | -0.90 | 0.37 |
|  | Opportunity Costs | 2.95 | 1.19 | 2.91 | 1.35 | -0.05<br>[-0.17, 0.07] | 455 | -0.82 | 0.41 |
| **Scientific Identity \*** |  | **3.89** | **1.08** | **4.19** | **1.19** | **0.29**<br>**[0.21, 0.38]** | **460** | **6.73** | **<0.001** |
| Intentions | Science Career | 4.41 | 0.84 | 4.42 | 0.84 | 0.01<br>[-0.05, 0.07] | 457 | 0.27 | 0.79 |
|  | Graduate Education | 3.99 | 1.00 | 3.96 | 1.11 | -0.04<br>[-0.12, 0.05] | 453 | -0.85 | 0.39 |
|  | Research Career | 3.16 | 0.99 | 3.11 | 1.08 | -0.04<br>[-0.12, 0.04] | 456 | -0.97 | 0.33 |
| Instructor Rapport |  | NA | NA | 5.61 | 1.17 | NA | NA | NA | NA |

**Table 4.** Effects of instructor talk on student outcomes. This table presents MLM results with standardized **ß and 95% confidence intervals**. ** **Bolded font** indicates p < 0.050 and * Indicates p < 0.10.

| Talk Type | Student Effect | $\beta$ (SE) | $df$ | $t$ | $p$ | CI |
| --- | --- | --- | --- | --- | --- | --- |
| Self-efficacy<br>Talk | Scientific Self-Efficacy | -0.01 (0.02) | 29 | -0.21 | 0.836 | [-0.14, 0.11] |
|  | Pursue graduate school | -0.05 (0.05) | 30 | -0.429 | 0.671 | [-0.29, 0.19] |
|  | Pursue a science career | -0.02 (0.04) | 27 | -0.142 | 0.8878 | [-0.27, 0.24] |
|  | Pursue a research career | 0.05 (0.04) | 25 | 0.479 | 0.636 | [-0.14, 0.23] |
| Task Values | Intrinsic Value * | 0.13 (0.05) | 38 | 1.75 | 0.090 | [-0.02, 0.28] |
|  | Attainment Value | 0.08 (0.04) | 35 | 1.24 | 0.224 | [-0.05, 0.20] |
|  | Social Utility * | 0.13 (0.04) | 30 | 1.93 | 0.063 | [0.00, 0.27] |
|  | Job Utility * | 0.12 (0.03) | 25 | 1.99 | 0.057 | [0.00, 0.25] |
|  | Life Utility | 0.08 (0.03) | 28 | 1.40 | 0.174 | [-0.03, 0.19] |
|  | Opportunity Costs | 0.008 (0.04) | 30 | 0.133 | 0.895 | [-0.12, 0.13] |
|  | **Scientific Identity \*\*** | **0.17 (0.04)** | **34** | **2.35** | **0.025** | **[0.03, 0.31]** |
|  | Pursue graduate school | 0.04 (0.04) | 33 | 0.67 | 0.51 | [-0.09, 0.17] |
|  | **Pursue a science career \*\*** | **0.18 (0.03)** | **31** | **2.57** | **0.015** | **[0.04, 0.32]** |
|  | Pursue a research career | 0.05 (0.03) | 33 | 1.01 | 0.322 | [-0.05, 0.16] |
| Immediacy | Student-Instructor Rapport * | 0.21 (0.07) | 39 | 1.85 | 0.072 | [-0.01, 0.43] |
|  | Pursue graduate school | -0.05 (0.06) | 30 | -0.474 | 0.639 | [-0.25, 0.15] |
|  | Pursue a science career | 0.003 (0.05) | 29 | 0.032 | 0.9751 | [-0.21, 0.21] |
|  | Pursue a research career | -0.01 (0.05) | 29 | -0.149 | 0.882 | [-0.17, 0.15] |

### Task values talk positively influences students’ scientific identity and marginally influences some but not all of students’ research task values

We hypothesized that instructors’ task values talk would boost students’ perceptions of the benefits of doing research, including its intrinsic value (i.e., how personally interesting or enjoyable doing research is), attainment value (i.e., how personally important doing research is), social utility (i.e., perceived value of research for making a difference in the world), job utility (i.e., perceived value of research for improving job prospects), and life utility (i.e., perceived value of research for daily life). We also hypothesized that instructors’ task values talk would reduce students’ perceptions of the opportunity costs of doing research (i.e., perception that doing research requires making sacrifices). Finally, because value beliefs shape a person’s self-concept and external scientific recognition (i.e., task values talk) foster a sense of belonging in science (Carlone & Johnson, 2007, 2007; Estrada et al., 2011; Pfeifer et al., 2024), we hypothesized that instructors’ task value talk would boost students’ scientific identity. To test these hypotheses and address our second research question (RQ2: *To what extent does instructor task values talk influence students’ post-course task value beliefs and scientific identity, controlling for pre-course levels?*), we conducted paired t-tests to compare students’ pre- and post-course task values and scientific identity. Consistent with prior research (Dunbar-Wallis et al., 2024; Hanauer et al., 2017; Newell & Ulrich, 2022), we observed that students reported significant increases in their scientific identity from pre- to post-course (Table 3). We observed no change in students’ task values except for their perceptions of the job utility of research, which dropped slightly but significantly from pre- to post-course.

In separate models for each student variable, we regressed students’ post-course task values and scientific identity onto the frequency of their instructor’s task values talk. We found that instructors’ task values talk had a significant but modest effect (ß=0.17, *p*=0.025) on their scientific identity (Table 4). This result suggests that instructors can favorably influence their students’ scientific identity when they are explicit about the nature of science, share their own interest or enthusiasm about science, provide career support, and comment on the larger purpose of students’ work (Zajic et al., 2026). We also found that instructors’ task values talk had small but marginally positive effects on students’ perceptions of the intrinsic value of research (ß=0.13, *p*=0.090), social utility of research (ß=0.13, *p*=0.063), and job utility of research (ß=0.12, *p*=0.057). These results suggest that instructors’ task value talk has the potential to influence students’ beliefs about the intrinsic value and social and job utility of research, but no effects on students’ beliefs about the attainment value, life utility, or opportunity costs of doing research. Notably, the overall trend in students’ perceptions of the job utility of research decreased significantly from pre- to post-course (i.e., negative effect), while instructors’ task values talk countered this effect (i.e., marginal positive effect).

### Immediacy talk is marginally associated with student ratings of rapport with their instructor

We hypothesized that students whose instructors used more immediacy talk, including sharing personal experiences, engaging in small talk, working to build trust, and emphasizing their availability (Zajic et al., 2026), would rate their rapport with their instructor more highly. To test this hypothesis and address our third research question (RQ3: *To what extent is instructor immediacy talk associated with student ratings of rapport with their instructor?*), we regressed students’ post-course ratings of instructor rapport onto the frequency of their instructor’s immediacy talk. We observed a marginal, positive association (ß=0.21, *p*=0.072) between student ratings of instructor rapport and instructors’ immediacy talk.

### The effect of instructor talk on students’ intentions is limited

Given our motivational framing, we hypothesized that students whose instructors used more self-efficacy, task values, and immediacy talk would report increased intentions to continue in science-related education and career paths. To test this hypothesis and address our final research question (RQ4: *To what extent do self-efficacy, task values, and immediacy talk influence students’ intentions to continue in science?*), we conducted paired t-test to compare students’ pre- and post-course intentions to pursue a science career, a science graduate degree, and a research career. We observed no differences. In separate models for each instructor talk-student outcome relationship, we regressed students’ post-course intentions onto the three types of instructor talk (i.e., nine models altogether). For the most part, instructor talk had no effect on students’ educational and career intentions. We observed one exception: instructors’ task value talk had a significant, positive effect on students’ intentions to pursue a science career (ß=0.18, *p*=0.015). This result suggests that instructors can favorably influence their students’ intentions to pursue a science career when they are explicit about the nature of science, share their own interest or enthusiasm about science, provide career support, and comment on the larger purpose of students’ work (Zajic et al., 2026). However, the effect of task values talk does not influence students’ research-related intentions, including pursuing a graduate degree or a research career.

## LIMITATIONS

Several aspects of our study limit what can be inferred. First, no prior research had quantitatively characterized the effects of non-content instructor talk on student-level variables. As a result, we could not draw from prior research to estimate effect sizes, and we were unable to calculate *a priori* the sample size needed to detect effects. Our results indicate that the effects we were able to detect were small (ß=0.12-21). Future research aimed at testing hypothesized relationships between instructor talk and student motivation should leverage our results to estimate sample sizes needed for robust tests. Second, we opted to use an MLM framework for our analysis instead of a structural equation modeling (SEM) framework. We made this decision to account for the nested nature of the data (students within instructors / courses / institutions) and because our sample size was insufficient for SEM. While we had a high number of second-level observations (i.e., 48 instructors/classes), many of these second-level observations contained less than ten first-level observations (i.e., student responses) rendering our sample insufficient for multi-level structural equation modeling (McNeish, 2017). Notably, MLM does not account for measurement error or allow for examination of potential paths between instructor talk, student beliefs, and student intentions. Finally, we fit multiple models to explore the effects of instructor talk on student outcomes, increasing the potential for Type 1 errors (false positives). Thus, our results should be interpreted with caution. Future research should examine whether the patterns we observed here are reproducible.

## DISCUSSION

Here we aimed to explore whether non-content instructor talk could play a motivating role in biology lab course instruction, possibly explaining the effects of CURE instruction on students’ continuation in college and in science (Corwin et al., 2018; Hanauer et al., 2017; Rodenbusch et al., 2016). We leveraged a previous EVT-informed characterization of instructor talk in 48 CURE and non-CURE courses to predict students’ motivation-related outcomes, including their scientific self-efficacy, task values, scientific identity, and science-related intentions. We were surprised that instructors’ self-efficacy talk had no observable effect on students’ scientific self-efficacy. Importantly, students reported increased scientific self-efficacy from pre- to post-course, thus some aspect of their experience is fostering their self-efficacy. Prior research indicates that mastery experience, meaning successful completion of a challenging task (Bandura, 1977), is a more influential source of self-efficacy than social persuasion, meaning what respected others say (Bandura, 1997; Capa-Aydin et al., 2018; Chen & Usher, 2013; Usher & Pajares, 2008). Our results are consistent with this – specifically that students are growing in their scientific self-efficacy because of what they are doing rather than what their instructors are saying. Recent research supports this interpretation, showing that, in comparison to other science practices students conduct during research, engaging in analytic tasks is a more influential source of scientific self-efficacy than engaging in other types of research-related tasks (e.g., experimentation, communication) (Zhang et al., 2026). It is important to note that the absence of an effect of self-efficacy talk should not be interpreted as permission to engage in negative forms of this talk. The qualitative analysis of instructor talk used as a foundation for this study revealed very few instances of “negative talk,” such as attributing experimental failures to students’ knowledge or skills or indicating that students should be more capable or competent. Thus, we were unable to test for any effects of negative self-efficacy talk. Research on undergraduate research experiences indicates that students whose research mentors engage in negative talk experience dampened self-efficacy and scientific identity growth as well as decreased intentions to continue in science (Limeri et al., 2024).

In contrast to self-efficacy talk, instructors’ task values talk had multiple positive effects on students’ motivation-related outcomes. Instructors’ task values talk favorably influenced students’ scientific identity. Instructors’ task values talk also had a marginally positive effect (*p*<0.057) on students’ beliefs about the value of doing research for their job prospects, countering a small but significant downward trend in their beliefs about the job utility of doing research from pre to post course. Furthermore, instructors’ task value talk also had positive effects on students’ beliefs about the intrinsic value and social utility of research, as well as their science-related career intentions. Collectively, these results suggest that instructors can favorably influence students’ value beliefs and persistence intentions by explaining what they find interesting or valuable about the work students are doing in class. This finding is consistent with the growing body of literature indicating that students opt out of science-related education and career paths not because of limited abilities but because of perceived misalignments with their values (Estrada et al., 2011; Jackson et al., 2016; Xia et al., 2024). Task values talk, which emphasizes the relevance or worth of students’ work or the course in general, may help students see connections between their schoolwork and what they personally value. Seeing this connection may help students see their beliefs as more aligned with the beliefs of the scientific community, fostering their identity as scientists and their intentions to continue in science.

On average, the CURE instructors in this study engaged in more task values talk than non-CURE instructors (Zajic et al., 2026), suggesting that CUREs may offer a curricular environment that is ripe for instructors to make the relevance and worth of students’ work more transparent. Indeed, this is consistent with early theory (Auchincloss et al., 2014; Corwin et al., 2015) and subsequent research (Cooper et al., 2019; Corwin et al., 2018) showing that CUREs are influential for students because they offer opportunities to make broadly relevant discoveries. However, it is important to note that task values talk is not limited to CUREs and CURE instructors vary widely in their use of task values talk (Zajic et al., 2026). This finding has practical implications because it suggests that any instructor – not just those who teach CUREs – could engage in task values talk to foster their students’ beliefs about the value of their coursework, their identification as scientists, and their intentions to pursue science-related careers. We encourage instructors to reflect on their curriculum and instruction to identify opportunities to explicitly reference how students’ tasks could be valuable beyond the course.

Future research should leverage our results to test the effects of task values talk more robustly, including a desired sample size selected based on the effect sizes we observed, multiple instructors within a course and institution to disaggregate effects of talk from other instructor, course, and institutional factors, and with more complete data from each instructor. Recent advances in large language models (LLMs) offer an avenue for replicating this work while reducing the labor required to do so. For example, every session from a lab course could be audio recorded, transcribed, and analyzed using an LLM training on the previous qualitative characterization of non-content talk. A larger sample would also enable analysis in an SEM framework, thus better accounting for measurement error. As noted previously, CURE instructors in our sample used more task values talk than non-CURE instructors, yet CURE instructors also varied widely in their task values talk (Zajic et al., 2026). A more robust assessment should take steps to ensure the “dose” of task values talk is consistent regardless of the course context. For instance, a future study could employ a randomized controlled design where instructors in the treatment condition use task values at some minimum frequency and instructors in the control condition do not. Brief instructor videos describing task values to students could help ensure consistent messaging and dose. Using a delayed-start design would also enable students in the control condition to benefit from task values talk.

Literature on the related constructs of verbal instructor immediacy (Christophel, 1990; Gorham, 1988; Mehrabian, 1966) and non-content instructor talk (Gelinas et al., 2022; Harrison et al., 2019; Ovid et al., 2021; Seidel et al., 2015) emphasizes their importance for building student-instructor relationships. We found that the relationship between instructors’ immediacy talk and students’ ratings of rapport with their instructor trended in a positive direction, but this relationship was not significant to *p*<0.05. It is important to note that the immediacy talk used to predict student ratings of rapport in this study does not reflect all of the categories of non-content instructor talk described previously (Gelinas et al., 2022; Harrison et al., 2019; Ovid et al., 2021; Seidel et al., 2015). This was intentional as we sought to link specific student outcomes with theoretically related types of talk. However, this may have missed other forms of non-content talk that could influence students’ rating of their rapport with their instructor or their overall perception of the classroom climate. Future research on non-content instructor talk could examine this by measuring additional forms of talk alongside theoretically related student experiences or outcomes.

Students in our study were in both CURE and non-CURE courses. The pre- to post-course differences we observed (e.g., gains in their scientific self-efficacy) were not limited to students in CUREs. In fact, the wide variation in talk we observed in the CUREs in our previous study motivated our decision to model our data using talk alone rather than course type. Collectively, our results show that forms of non-content talk can be influential for students, but this influence is not apparent in CUREs alone or in all CUREs.

This finding adds to an emerging pattern in the literature that CUREs – or other types of undergraduate research or laboratory courses – do not constitute a singular experience (Beck et al., 2023; Dolan, 2016; Gentile et al., 2017; Zhang et al., 2026). Rather, future research on CUREs and other forms of practical instruction must examine how features of their design and implementation relate to students’ experiences and outcomes. Results of this research are needed to understand how CUREs are influential for students and to inform the design and implementation of CUREs.

## Supporting information

Zajic and Dolan Supplemental Materials

## ACKNOWLEDGMENTS

We are grateful to our participants for allowing us to record their instruction and to their students for their survey responses. We are also grateful to Benjamin Listyg and Christina Leckfor for their guidance on data cleaning and statistical analyses as well as Rob Erdmann, Paul Hernandez, and Emily Rosenzweig for their feedback on the analyses. We appreciate Alexandra Cooper, Trevor Tuma, and other members of the Social Psychology of Research Experiences and Education (SPREE) research group and the Biology Education Research Group for feedback on work in progress and drafts of this manuscript. Finally, we would like to thank Tessa Andrews and Jennifer Jo Thompson for their thoughtful feedback. This work was supported in part by National Science Foundation Division of Undergraduate Education Improving Undergraduate STEM Education Collaborative Grant (Awards 2021138 and 2021112), an NSF Graduate Research Fellowship (Award 2236869), and the Georgia Athletic Association Professorship for Innovative Science Education. Any opinions, findings, conclusions, or recommendations expressed in this material are those of the authors and do not necessarily reflect the views of any of the funding organizations.

