## Supplementary material for "Influence of non-content instructor talk on students’ motivation-related outcomes in laboratory courses": Zajic and Dolan Supplemental Materials

This supplement contains the following:

| Item | Page |
|---|---|
| Measure Descriptions, Descriptive Stats, and Measurement Models | S2-S8 |
| Tables S1A-C. Scientific self-efficacy | S2 |
| Tables S2A-C. Science identity | S3 |
| Tables S3A-C. Task value – benefits | S4-5 |
| Tables S4A-C. Task value – opportunity cost | S6 |
| Table S5A-C. Instructor rapport | S7 |
| Tables S6-S8. Intentions | S8 |
| Figure S1. Correlation matrix | S9 |
| Table S6. Intraclass correlations | S10 |

### MEASURE DESCRIPTIONS, DESCRIPTIVE STATISTICS, AND MEASUREMENT MODELS

**Construct:** Scientific Self-Efficacy
**Instructions:** Please indicate how confident you are in your ability to...
**Response Format:** Not confident = 1, A little confident = 2, Somewhat confident = 3, Confident = 4, Very Confident = 5, Extremely Confident = 6, I prefer not to respond = NA

**Table S1A. Descriptive statistics for scientific self-efficacy responses**

| Variable | Range | Mean (Standard Deviation) |
|---|---|---|
| Scientific Self-Efficacy (Pre) | 2-6 | 3.58 (0.81) |
| Scientific Self-Efficacy (Post) | 1.56-6 | 4.19 (0.94) |

**Table S1B. Item descriptions and factor loadings for measure of scientific self-efficacy**

| Item | Content | Pre | Post |
|---|---|---|---|
| SSE_1 | Use technical skills (lab or field equipment, instruments, and/or bench or field techniques). | 0.58 | 0.56 |
| SSE_2 | Use computational skills (software, algorithms, and/or quantitative techniques). | 0.47 | 0.54 |
| SSE_3 | Generate a research question to answer. | 0.75 | 0.87 |
| SSE_4 | Develop a hypothesis to test. | 0.77 | 0.87 |
| SSE_5 | Figure out what data/observations to collect and how to collect them. | 0.80 | 0.82 |
| SSE_6 | Trouble-shoot an investigation or experiment. | 0.71 | 0.76 |
| SSE_7 | Create explanations for the results of the study. | 0.72 | 0.74 |
| SSE_8 | Use scientific literature and/or reports to guide research | 0.65 | 0.67 |
| SSE_9 | Develop theories (integrate and coordinate results from multiple studies). | 0.72 | 0.76 |

**Table S1C. Confirmatory factor model fit statistics for scientific self-efficacy scale***

| Model | CFI | TLI | RMSEA | SRMR | $\chi 2$ | *df* |
|---|---|---|---|---|---|---|
| Scientific Self-Efficacy Pre | 0.861 | 0.815 | 0.151 | 0.06 | 278 | 27 |
| Scientific Self-Efficacy Post | 0.906 | 0.875 | 0.139 | 0.04 | 265 | 27 |

*** Note.** Typical cutoff values for global fit statistics indicative of acceptable model fit are a comparative fit index (CFI) $\geq 0.95$, Tucker-Lewis fit index (TLI) $\geq 0.95$, root mean squared error of approximation (RMSEA) $\leq 0.06$, and standardized root mean squared residual (SRMR) $\leq 0.08$. $\chi 2$ = the chi-squared test statistic for the model. df = degrees of freedom.

**Construct:** Science Identity

**Instructions:** Please indicate the extent to which you agree with the following statements.

**Response Format:** Strongly disagree = 1, Moderately disagree = 2, Slightly agree = 3, Moderately agree = 4, Mostly agree = 5, Strongly agree = 6, I prefer not to respond = NA

**Table 2A. Descriptive statistics for science identity responses**

| Variable | Range | Mean (Standard Deviation) |
|---|---|---|
| Science Identity (Pre) | 1-6 | 3.89 (1.08) |
| Science Identity (Post) | 1-6 | 4.19 (1.19) |

**Table 2B. Item descriptions and factor loadings for measure of science identity**

| Item | Content | Pre | Post |
|---|---|---|---|
| SI_1 | I have a strong sense of belonging to the community of scientists. | 0.63 | 0.76 |
| SI_2 | I derive great personal satisfaction from working on a team of scientists. | 0.66 | 0.74 |
| SI_3 | I think of myself as a scientist. | 0.72 | 0.77 |
| SI_4 | The daily work of a scientist is appealing to me. | 0.63 | 0.77 |
| SI_5 | I feel like I belong in the field of science. | 0.65 | 0.71 |
| SI_6 | In general, being a scientist is an important part of my self-image. | 0.90 | 0.91 |
| SI_7 | Being a scientist is an important reflection of who I am. | 0.91 | 0.91 |

**Table 2C. Confirmatory factor model fit statistics for science identity scale**

| Model | CFI | TLI | RMSEA | SRMR | $\chi 2$ | *df* |
|---|---|---|---|---|---|---|
| Science Identity Pre | 0.867 | 0.800 | 0.200 | 0.07 | 268 | 14 |
| Science Identity Post | 0.897 | 0.845 | 0.199 | 0.05 | 269 | 14 |

**Construct:** Benefits

**Instructions:** Please indicate the extent to which you agree with the following statements.

**Response Format:** Strongly disagree = 1, Moderately disagree = 2, Slightly agree = 3, Moderately agree = 4, Mostly agree = 5, Strongly agree = 6, I prefer not to respond = NA

**Table 3A. Descriptive statistics for beneficial task values responses**

| Variable | Range | Mean (Standard Deviation) |
|---|---|---|
| Intrinsic Value (Pre) | 1-6 | 4.33 (1.23) |
| Intrinsic Value (Post) | 1-6 | 4.40 (1.36) |
| Attainment Value (Pre) | 1.33-6 | 4.78 (1.12) |
| Attainment Value (Post) | 1-6 | 4.70 (1.28) |
| Social Utility (Pre) | 1-6 | 4.81 (1.00) |
| Social Utility (Post) | 1-6 | 4.74 (1.07) |
| Job Utility (Pre) | 1-6 | 5.18 (0.91) |
| Job Utility (Post) | 1-6 | 5.02 (1.02) |
| Life Utility (Pre) | 1.33-6 | 4.55 (1.03) |
| Life Utility (Post) | 1-6 | 4.51 (1.16) |

**Table 3B. Item descriptions and factor loadings for measure of research task values**

| Item | Content | Pre | Post |
|---|---|---|---|
| **Intrinsic Value** | | | |
| B_1 | Research is fun to me. | 0.95 | 0.96 |
| B_2 | I like doing research. | 0.97 | 0.97 |
| B_3 | I enjoy dealing with research topics. | 0.90 | 0.90 |
| **Attainment Value** | | | |
| B_4 | It is important to me to be good at research. | 0.88 | 0.89 |
| B_5 | Being good at research means a lot to me. | 0.91 | 0.95 |
| B_6 | Performing well in research is important to me. | 0.85 | 0.90 |
| **Social Utility** | | | |
| B_7 | Being well versed in research will prepare me to help my community. | 0.79 | 0.83 |
| B_8 | I can do good in the world based on my knowledge of research. | 0.75 | 0.87 |
| B_9 | If I know a lot about research, I can make a difference in the world. | 0.74 | 0.81 |
| **Job Utility** | | | |
| B_10 | Doing well in research will improve my chances of finding a job after college. | 0.81 | 0.86 |
| B_11 | The skills I develop in research will help me be successful in my career. | 0.85 | 0.86 |
| B_12 | Learning how to conduct research is worthwhile because it improves my career prospects. | 0.83 | 0.89 |
| **Life Utility** | | | |
| B_13 | Research will help me in life. | 0.88 | 0.90 |

| | | | |
|---|---|---|---|
| B_14 | I will often need research in my life. | 0.77 | 0.83 |
| B_15 | Research comes in handy in everyday life. | 0.69 | 0.73 |

**Table 3C. Confirmatory factor model fit statistics for benefits scales**

| Model | CFI | TLI | RMSEA | SRMR | $\chi 2$ | *df* |
|---|---|---|---|---|---|---|
| Benefits Pre | 0.984 | 0.980 | 0.047 | 0.03 | 159 | 80 |
| Benefits Post | 0.979 | 0.973 | 0.060 | 0.03 | 212 | 80 |

**Construct:** Opportunity Cost

**Instructions:** Please indicate the extent to which you agree with the following statements.

**Response Format:** Strongly disagree = 1, Moderately disagree = 2, Slightly agree = 3, Moderately agree = 4, Mostly agree = 5, Strongly agree = 6, I prefer not to respond = NA

**Table 4A. Descriptive statistics for opportunity cost responses**

| Variable | Range | Mean (Standard Deviation) |
|---|---|---|
| Opportunity Cost (Pre) | 1-6 | 2.95 (1.19) |
| Opportunity Cost (Post) | 1-6 | 2.91 (1.35) |

**Table 4B. Item descriptions and factor loadings for measure of opportunity costs associated with participation in research**

| Item | Content | Pre | Post |
|---|---|---|---|
| OC_1 | I have to give up other activities that I like to be successful at research. | 0.63 | 0.76 |
| OC_2 | I have to give up a lot to do well in research. | 0.66 | 0.74 |
| OC_3 | I'd have to sacrifice a lot of free time to be good at research. | 0.72 | 0.77 |

**Table 4C. Confirmatory factor model fit statistics for opportunity cost scale**

| Model | CFI | TLI | RMSEA | SRMR | $\chi 2$ | *df* |
|---|---|---|---|---|---|---|
| Opportunity Cost (Pre) | 1.000 | 1.000 | 0.000 | 0.00 | 0 | 0 |
| Opportunity Cost (Post) | 1.000 | 1.000 | 0.000 | 0.00 | 0 | 0 |

**Construct:** Instructor Rapport

**Instructions:** Please indicate the extent to which each description is indicative of your instructor.

**Response Format:** Strongly disagree = 1, Disagree = 2, Somewhat disagree = 3, Neither agree nor disagree = 4, Somewhat agree = 5, Agree = 6, Strongly agree = 7, I prefer not to respond = NA

**Table 5A. Descriptive statistics for instructor rapport responses**

| Variable | Range | Mean (Standard Deviation) |
|---|---|---|
| Instructor Rapport (Post) | 1-7 | 5.61 (1.17) |

**Table 5B. Item descriptions and factor loadings for measure of instructor rapport**

| Item | Content | Post |
|---|---|---|
| IR_1 | I enjoy interacting with my instructor. | 0.87 |
| IR_2 | My instructor creates a feeling of "warmth" in our relationship. | 0.87 |
| IR_3 | My instructor relates well to me. | 0.87 |
| IR_4 | I have a harmonious relationship with my instructor. | 0.84 |
| IR_5 | My instructor has a good sense of humor. | 0.74 |
| IR_6 | I am comfortable interacting with my instructor. | 0.83 |
| IR_7 | I feel like there is a "bond" between my instructor and myself. | 0.87 |
| IR_8 | I look forward to seeing my instructor in class. | 0.92 |
| IR_9 | I strongly care about my instructor. | 0.80 |
| IR_10 | My instructor has taken a personal interest in me. | 0.72 |
| IR_11 | I have a close relationship with my instructor. | 0.73 |

**Table 5C. Confirmatory factor model fit statistics for instructor rapport scale**

| Model | CFI | TLI | RMSEA | SRMR | $\chi 2$ | *df* |
|---|---|---|---|---|---|---|
| Instructor Rapport Post | 0.865 | 0.831 | 0.186 | 0.06 | 743 | 44 |

**Single Item Measures**

**Construct:** Graduate School Intentions

**Instructions:** To what extent do you intend to pursue a graduate degree in science?

**Response Format:** I DEFINITELY WILL NOT pursue a graduate degree in science = 1, I PROBABLY WILL NOT pursue a graduate degree in science = 2, I am UNSURE whether I will pursue a graduate degree in science = 3, I PROBABLY WILL pursue a graduate degree in science = 4, I DEFINITELY WILL pursue a graduate degree in science = 5, I prefer not to respond = NA

**Table S6. Descriptive statistics for graduate school intentions responses**

| Variable | Range | Mean (Standard Deviation) |
|---|---|---|
| Graduate School Intentions (Pre) | 1-5 | 3.99 (1.00) |
| Graduate School Intentions (Post) | 1-5 | 3.96 (1.11) |

**Construct:** Science Career Intentions

**Instructions:** To what extent do you intend to pursue a career in science?

**Response Format:** I DEFINITELY WILL NOT pursue a career in science = 1, I PROBABLY WILL NOT pursue a career in science = 2, I am UNSURE whether I will pursue a career in science = 3, I PROBABLY WILL pursue a career in science = 4, I DEFINITELY WILL pursue a career in science = 5, I prefer not to respond = NA

**Table S7. Descriptive statistics for science career intentions responses**

| Variable | Range | Mean (Standard Deviation) |
|---|---|---|
| Science Career Intentions (Pre) | 1-5 | 4.41 (0.84) |
| Science Career Intentions (Post) | 1-5 | 4.42 (0.87) |

**Construct:** Science Research Career Intentions

**Instructions:** To what extent do you intend to pursue a science research-related career?

**Response Format:** I DEFINITELY WILL NOT pursue a science research-related career = 1, I PROBABLY WILL NOT pursue a science research-related career = 2, I am UNSURE whether I will pursue a science research-related career = 3, I PROBABLY WILL pursue a science research-related career = 4, I DEFINITELY WILL pursue a science research-related career = 5, I prefer not to respond = NA

**Table S8. Descriptive statistics for science research career intentions responses**

| Variable | Range | Mean (Standard Deviation) |
|---|---|---|
| Science Research Career Intentions (Pre) | 1-5 | 3.16 (0.99) |
| Science Research Career Intentions (Post) | 1-5 | 3.11 (1.08) |

**Figure S1. Correlations.** This figure displays the correlation matrix for all pre and post student-level variables.

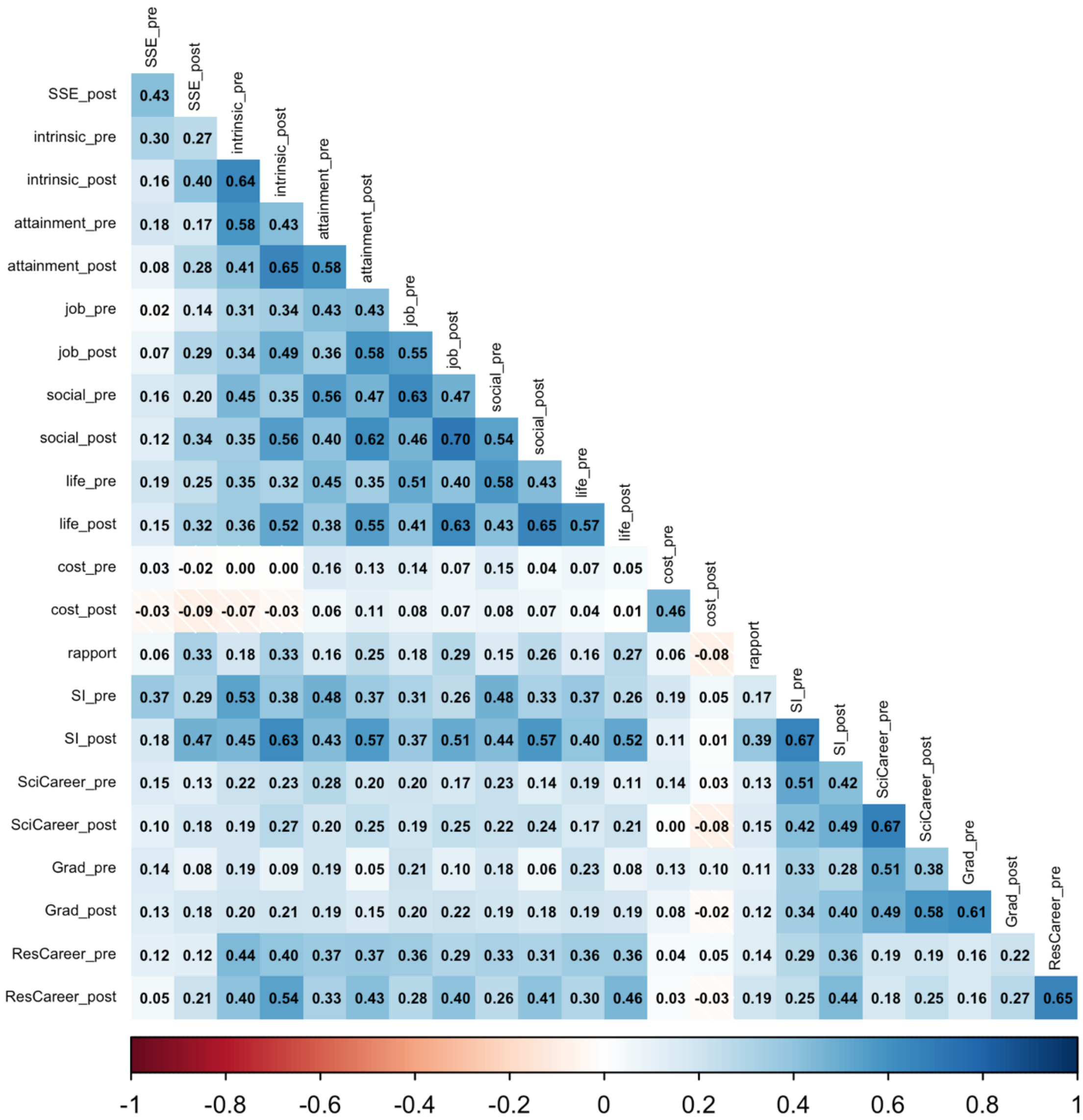


**Label Key:**

SSE = Scientific Self-Efficacy

SI = Science Identity

Grad = Graduate School in Science Intentions

SciCareer = Career in Science Intentions

ResCareer = Research Career in Science Intentions

*The words "utility" and "value" are left out of the task value related variables (e.g., "intrinsic_pre" = the pre-course score on Intrinsic Value items)

**Table S9. Intraclass Correlations.** Here we report the within and between-group variances for each measured student outcome measured at the post timepoint and grouped by instructor.

| Outcome | Between-Group Variance | Within-Group Variance |
|---|---|---|
| Scientific Self-Efficacy | 3.9% | 96.1% |
| Science Identity | 11.5% | 88.5% |
| Intrinsic Value | 15.4% | 84.6% |
| Attainment Value | 6.8% | 93.2% |
| Social Utility | 9.6% | 90.4% |
| Job Utility | 7% | 93% |
| Life Utility | 3% | 97% |
| Opportunity Costs | 7.1% | 92.9% |
| Graduate School Intentions | 6.5% | 93.5% |
| Science Career Intentions | 11.2% | 88.8% |
| Research Career Intentions | 1% | 99% |
| Rapport with Instructor | 38.7% | 61.3% |
